# Primitive Molecular Buffering by Low-Multivalency Coacervates

**DOI:** 10.1101/2025.05.22.655649

**Authors:** Saehyun Choi, Sindy P. Liu, McCauley O. Meyer, Philip C. Bevilacqua, Christine D. Keating

## Abstract

Coacervate droplets formed by liquid-liquid phase separation serve as models for intracellular biomolecular condensates and as potential protocellular compartments during the emergence of life. Changes in availability of molecular components can be anticipated for intracellular and prebiotic milieu, and protocells may have also faced fluctuations in salinity and pH. Compartments able to maintain their molecular composition, i.e. homeostasis, under such conditions would be better able to preserve internal functions. Phase separation could in principle provide resistance to local changes in molecular composition. To evaluate this possibility, we investigated the impact of non-stoichiometric charge ratios of coacervate molecules on coacervate formation and RNA compartmentalization in oligoarginine (R10)/ATP coacervates across salinity and pH conditions relatable to plausible prebiotic environments. These R10/ATP coacervate systems resisted changes in oligoarginine concentration in both phases under freshwater and ocean-relevant salt conditions, providing a primitive molecular buffering function. Moreover, RNA accumulation was observed in the coacervates over a range of pH, salinity, and R10/ATP stoichiometry. We also observed salt-dependent differences in molecular buffering and compartmentalization that can be understood in terms of how salinity impacts the relative strengths of intermolecular binding modes that drive coacervation and RNA uptake. By varying relative phase volumes and altering which intermolecular binding modes dominate, LLPS provides general mechanisms for resisting changes in molecular availability and environmental conditions, even without the active homeostasis of living cells. Such primitive molecular buffering could have aided the emergence of life and may find utility in biotechnological or commercial applications based on molecular compartmentalization.

## Introduction

Coacervate droplets form by liquid-liquid phase separation (LLPS) and serve as leading model systems for prebiotic compartments^1^ and membraneless biological condensates in cells^2^. These structures have locally high concentrations of their component molecules and a remarkable ability to localize guest molecules ranging from small molecules to biomacromolecules.^1, 3^ Indeed, accumulation of relatively rare and otherwise dilute (proto)-biomolecules from lakes, oceans, or other prebiotic waters could have facilitated the emergence of life.^1, 4, 5^ The LLPS-generated microenvironments of membraneless organelles, which contain intrinsically disordered proteins (IDPs) and often RNA, support numerous functions in extant biology including RNA sequestration and processing,^6–8^ suggesting the possibility that prebiotic coacervates may have offered similar capabilities. In particular, encapsulation of RNA or RNA-related molecules in coacervates could aid scenarios relating to RNA world^9, 10^ or RNA/peptide world^11, 12^ hypotheses by increasing local RNA concentrations, providing protection from degradation, influencing RNA-RNA and RNA-peptide interactions, and increasing ribozyme kinetics.^1, 13–16^ Molecular components of prebiotic coacervates would presumably have been less complex and lower in molecular weight than the proteins and nucleic acid polymers of today’s membraneless organelles. Although simpler and shorter molecules, including homopeptides of ten or fewer residues, can undergo LLPS and the resulting coacervates can perform well as RNA compartments,^17, 18^ their lower-multivalency is expected to render them more susceptible to dissolution and less able to maintain distinct internal microenvironments as compared to coacervates formed from longer polymers^19^. Compounding potential concerns, prebiotic waters are thought to include a wide range of salinity^20^ and pH^21^ conditions that vary from groundwater to seawater^21, 22^ and include pH gradients associated with geochemical features such as hydrothermal vents^23^ upon which coacervation depends.^2, 18, 24^ It is thus critical to assess the impact of such environmental conditions on compartmentalization functions such as guest molecule uptake.

Maintaining steady concentrations of intracellular biomolecules is an important aspect of homeostasis in extant biology. Coacervates formed by LLPS might also resist changes in concentrations of phase-forming components to provide a primitive molecular “buffering” mechanism. For example in simple phase-separating systems where a polymer self-associates to form polymer-rich and -poor phases, changes in the availability of the polymer can be accommodated by changes in phase volumes, such that the compositions of both phases remain unchanged.^25, 26^ Indeed, it has been hypothesized that membraneless organelles of extant cells may protect against local intracellular protein concentration fluctuations and thus from expression noise in this way,^27, 28^ and similar molecular buffering might be expected for simpler protocellular systems. However, the effectiveness of phase separation in maintaining local biomolecule concentrations in multicomponent LLPS systems is less straightforward and not yet well understood.^29, 30^ Even for relatively simple two-component LLPS systems such as complex coacervates in which one polycation and one polyanion are mixed, molecular buffering expectations are not clear, particularly when relative amounts of the two components (e.g., the ratio of cationic to anionic functional groups) are varied. Changes in the overall cationic/anionic charge ratio for a two-component system can be expected to require a change in the composition of at least one of the phases. In the simplest case, coacervate composition could remain unchanged, accommodated by the “excess” polyion accumulating in the dilute phase and/or coacervate volume changes in response to the availability of a limiting polyion. In this scenario, concentrations of both polyions are well-buffered in the coacervate phase, and the concentration of one polyion –but not both– could also exhibit a resistance to change in the dilute phase. Alternatively, the compositions of both the coacervate and dilute phases could change with cationic/anionic charge ratio; variations in relative phase volumes are also likely in this scenario.^31, 32^ The extent to which either phase would exhibit molecular buffering for either polyion in these systems is harder to predict. All scenarios where the overall ratio of coacervate-forming components is altered can be expected to impact the distribution of any guest molecules present in the system, potentially impacting the performance of coacervate droplets as compartments.

When considering the ability of phase separation to provide molecular buffering, it is essential to consider what molecules and what phases are most important. For example, in living cells, where the outside and inside of the droplet are both part of the cell, it may be more important for expression noise filtering to avoid protein concentration fluctuations outside than inside the droplet phase, while in prebiotic compartmentalization scenarios it may be more important to maintain homeostasis in the droplet interior since in the simplest scenario it comprises the entire protocell. Because phase volume changes provide the most straightforward mechanism for buffering of coacervate-forming molecules, situations leading to optimal buffering of the coacervate-forming molecules (by growing or shrinking the coacervate phase volume) may be incompatible with simultaneously maintaining local guest solute concentrations within the compartment according to changes in encapsulation fraction of guest molecules (by the dilution or concentration, respectively, of guest molecules as compartment volume changes).

Here, we evaluated whether coacervates formed from relatively simple, low molecular-weight molecular components perform effectively as RNA compartments when challenged by nonstoichiometric cationic/anionic moiety mixing ratios and environmental changes (e.g. simulating pH gradient in hydrothermal vents^23^ or seawater versus groundwater^22^). We report that complex coacervates formed by mixing cationic oligoarginine (R10) and anionic ATP are stable and function as model prebiotic compartments under both low and high ionic strength as well as weakly basic to acidic pH, even in the presence of changes in the relative amounts of R10 vs ATP. We find that the local composition of R10/ATP coacervates is less impacted by changes in R10/ATP ratio at low salt and high pH condition, while ATP and RNA concentrations within the coacervates are significantly enhanced in high salt condition. These seemingly counterintuitive results can be understood as resulting from how ionic strength and pH impact the relative strengths of R10-ATP interaction modes. Even without the active homeostasis of living cells, this two-component phase-separating system resisted changes in phase composition and the microenvironment experienced by RNA guest molecules, supporting the potential of multicomponent LLPS as a primitive molecular buffering mechanism.

## Results and Discussion

We chose to investigate coacervates formed from R10 and ATP over a range of cationic/anionic moiety charge ratios. Both polyions are relatively low-multivalency, which is relevant to early Earth conditions; moreover, both components are important in extant biology. Arginine, in particular, is critical for protein phase separation^33–35^ and interactions with nucleic acids via ionic pairing, cation-*π* and *π* – *π* interactions.^33, 36 37^ While arginine is not thought to be one of the very earliest amino acids formed, prebiotically-plausible syntheses have been proposed^38, 39^, and the lower reactivity of arginine’s guanidinium moieties (as compared for example to lysine’s primary amines) would have aided its persistence. Extant cells have high levels of ATP, between approximately 0.5 and 10 mM depending on cell type,^40^ which have been suggested to play important roles in intracellular LLPS.^41–44^ Moreover, ATP can be produced by prebiotically-plausible synthetic routes.^45–47^ The high salt stability of R10/ATP coacervation (critical salt concentration ∼700 mM KCl^18^) makes it possible to evaluate these coacervates over ionic strengths that resemble both freshwater^20, 21, 48^ (15 mM KCl, low salt condition) and oceanic salt conditions (600 mM KCl, high salt condition). In our experiments, both conditions have 0.5 mM MgCl_2_ for RNA duplex formation. We began by examining changes in R10 and ATP concentration in the coacervate and dilute phases at different ratios and ionic strengths.

### Molecular buffering by coacervation

Equal-charge (1:1) matching of cationic and anionic moieties, which is favorable for LLPS, is typically used in laboratory studies of complex coacervates, but cannot be assumed in realistic prebiotic or biological scenarios. This led us to ask whether LLPS still occurs in solutions of non-stoichiometric charge ratios for relatively low molecular weight molecules and whether any resulting coacervates still retain their roles as compartments. Reasoning that ATP might be more prevalent than arginine peptides prebiotically due to ATP’s smaller molecular weight and higher relative abundance in prebiotic syntheses^45^, we chose to fix the ATP concentration at 10 mM, while varying the R10 concentration from 0.5 – 4.0 mM. These experiments were performed at pH 8.1 ∼ 8.5, which fully deprotonates the triphosphate and is considered relevant to the pH of spring water and oceans; moreover, it allows us to compare with our prior work^18^ at stoichiometric charge ratio and decalysine (K10)/ATP coacervates in low salt condition at pH 8.1 ∼ 8.5. We will refer to concentration of molecules in each phase as [MOI]_p,_ where ‘MOI’ indicates the molecule of interest and ‘p’ indicates the phase of interest. The unit ‘mM_c_’ will be used to refer to the charge concentration, in mM, associated with a particular polyion. Since ATP is assumed to be fully deprotonated at pH 8.1 ∼ 8.5, 10 mM [ATP]_total_ has a charge concentration of 40 mM_c_. Varying [R10]_total_ from 0.5 –4.0 mM corresponds to charge concentrations ranging from 5 to 40 mM_c_ , with R10/ATP charge ratios spanning from 0.5/4 to 4/4. We confirmed coacervate formation under this range of stoichiometries in both low and high salt conditions (see **Figure S1**). We then characterized the R10 and ATP content of the coacervate and dilute phases as a function of R10/ATP ratio under low and high salt conditions. Concentrations of R10 were determined using TAMRA-labeled R10 (**Figure 1A**), while concentrations of ATP were determined by UV absorbance at 260 nm (**Figure 1B**), as described in Methods and **Figure S2**.

**Figure 1.**
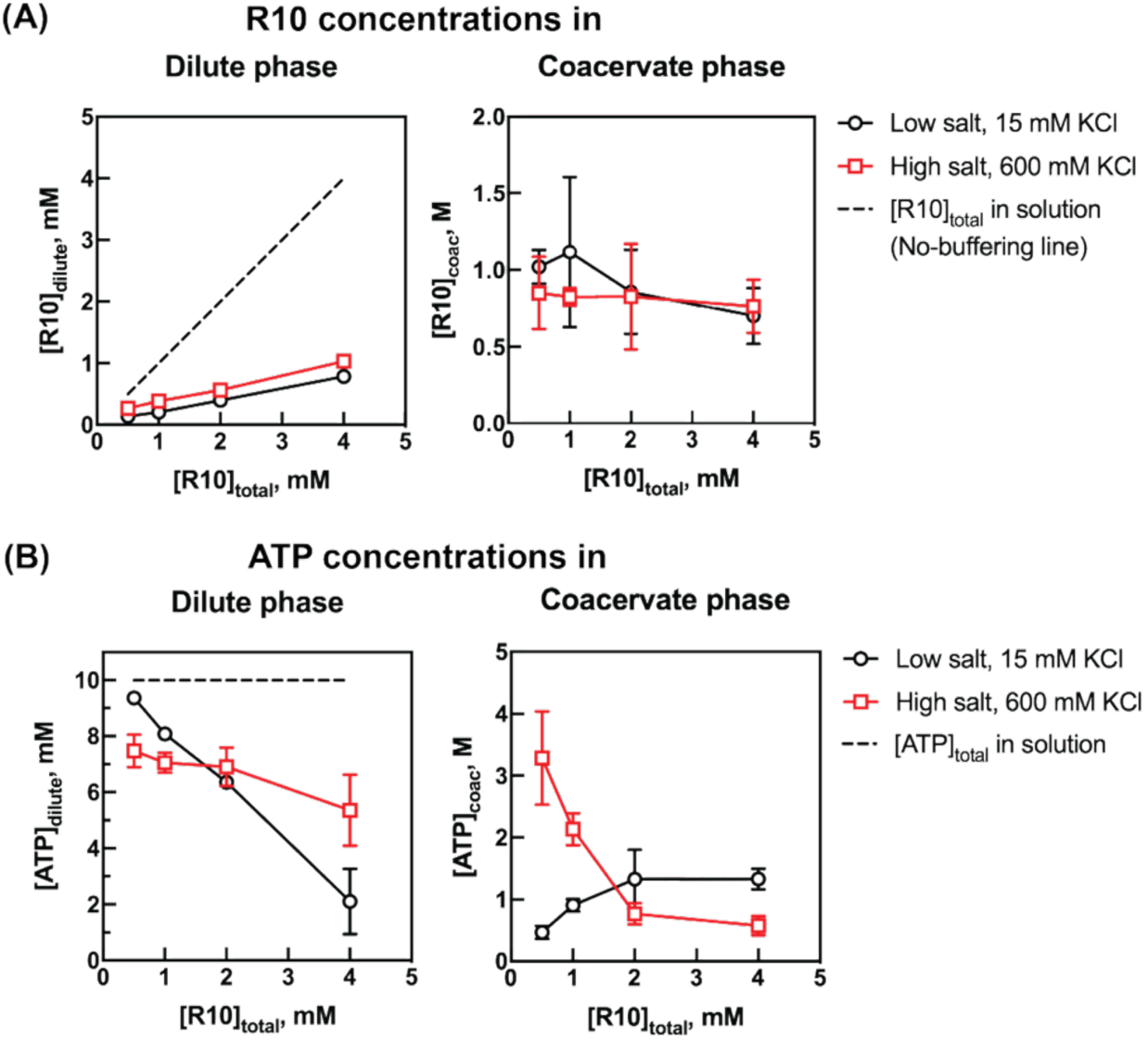
Quantification of R10 and ATP in coacervate phase and dilute phase of different charge ratios. **(A)** R10 concentrations in dilute and coacervate phases and (**B**) ATP concentrations in dilute and coacervate phases. X-axis denotes total R10 concentration added, not accounting for phase separation, while total ATP concentrations are all fixed at 10 mM. All conditions are pH 8.5 and contain 0.5 mM MgCl_2_. Data points indicate the average values, and error bars indicate the standard deviation of three independent trials. Average, standard deviation, and statistical analysis results (t-tests, Pearson’s correlation test, Spearman’s rank test) are provided in **Table S1-S3.**

We first asked how R10 concentrations in coacervate phase and dilute phases respond to [R10]_total_ changes: would either phase show buffering of [R10]? When [R10]_total_ was increased 8-fold from 0.5 to 4 mM (5 to 40 mM_c_) at a fixed [ATP]_total_ of 10 mM (40 mM_c_), we found relatively little of the added R10 in the dilute phase (**Figure 1A**, left panel). [R10]_dilute_ increased roughly proportionately to [R10]_total_, from 0.1 to 0.8 mM for low salt condition and from 0.3 to 1.0 mM for high salt condition, indicating the rest of the added R10 must have been accommodated by the coacervate phase. Turning to the coacervate phase, the [R10]_coac_, was estimated to be between ∼1 and 0.8 M for low and high salt samples, or ∼1,000 to 10,000 times greater than [R10]_dilute_, independent of the initial mixing charge ratios of R10 and ATP (**Figure 1A**, right panel). The steepness of the slope in **Figure 1A** indicates how R10 concentrations vary in the dilute and coacervates, as bulk concentrations of R10 are changed; this slope defines molecular buffering capacity, with shallow or zero slopes indicating stronger buffering. For both low and high salt concentrations, the slope of [R10]_dilute_ vs. [R10]_total_ was 0.2 from linear regression, which is much less than the no-buffering line of 1, indicating strong resistance of changes in [R10]_dilute_ to changes in [R10]_total_, or potent buffering of R10 in the dilute phase (**Figure 1A**; [R10] values are also provided in **Table S1**). The slope of [R10]_coac_ vs. [R10]_total_ was near 0, indicating nearly complete resistance of changes in [R10]_coac_ to changes in [R10]_total_, or nearly complete buffering in the coacervate phase. In other words, the protocell portion of the system is exceptionally resistant to change in local [R10].

The observation that R10/ATP coacervate systems resist changes in [R10]_dilute_ and [R10]_coac_ with increasing [R10]_total_ suggested that the added R10 molecules are accommodated in the coacervate by forming increasing the coacervate phase volume. Indeed, inspection of centrifuged samples clearly shows larger coacervate phase volumes with increasing [R10]_total_ (**Figure 2**), supporting the formation of additional coacervate phase as a primary mechanism by which the coacervate system buffers the local R10 concentrations ([R10]_coac_, [R10]_dilute_) even as the total amount of R10 in the system ([R10]_total_) increases. If the R10:ATP stoichiometry of the coacervate phase is maintained as it grows, then [ATP]_dilute_ can be expected to decrease with increasing [R10]_total,_ to provide the ATP necessary to grow the coacervate. Indeed, we found that [ATP]_dilute_ decreased with increasing [R10]_total_ (**Figure 1B, Table S2)**. This trend was more pronounced for the low salt condition. From linear regression, the slopes of [ATP]_dilute_ vs. [R10]_total_ were -2 and -0.5 for low salt and high salt condition, respectively. This indicates differences in the degree to which these two systems change [ATP] to maintain their coacervate phase composition when faced with added [R10]_total_, which may in part reflect the greater availability of chloride anions to perform part of the charge balance in the high salt samples resulting in less depletion of ATP from the bulk.

**Figure 2.**
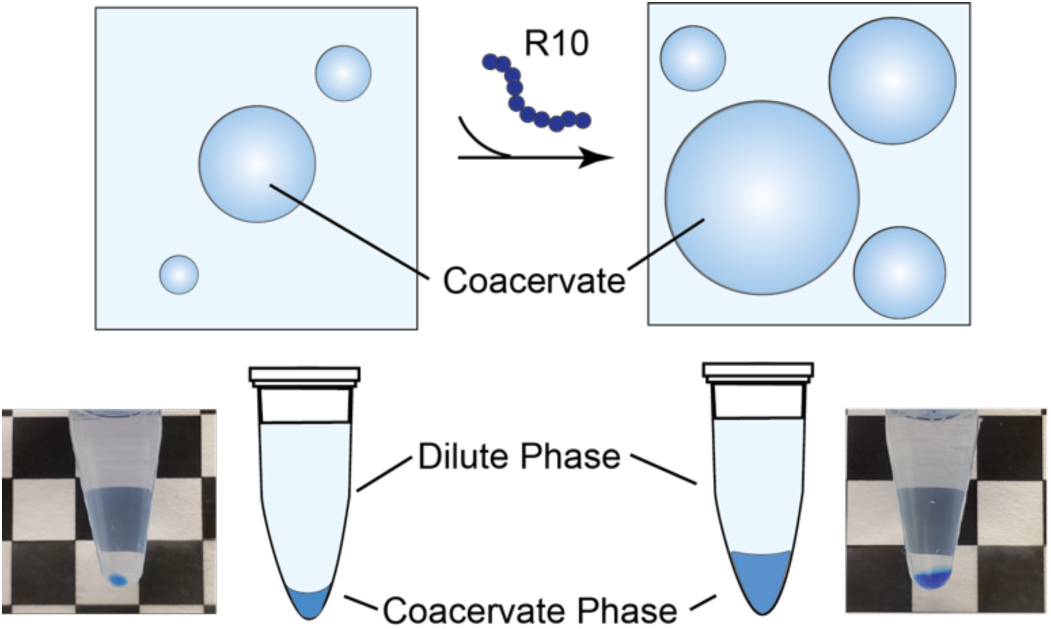
Coacervate phase volume increases upon addition of R10. The left and right side-view images show coacervate samples with 1 and 4 mM [R10]_total_ under low salt conditions, respectively. The coacervate phase is visualized by adding 15 µM of bromothymol blue. The remaining coacervate phase images at different [R10]_total_ conditions are provided in Figure S3.

Notably, the [ATP]_coac_ trends are opposite for the two salt conditions. To determine [ATP]_coac_, we used values for [ATP]_total_ and [ATP]_dilute_ and an approximated volume for the coacervate phase (V_coac_). Direct measurement of coacervate phase volume is challenging at current peptide and ATP concentrations.^49, 50^ Therefore, we estimated the coacervate volume in two different ways: from the zoomed-in coacervate phase images (**Figure S3(A)-(D)**) and from R10 concentrations (**Figure S3E**) (Methods). The two methods agreed within a factor of 2 (**Figure S3E**). Gratifyingly, these independent approaches to coacervate phase volume estimation gave similar trends of [ATP]_coac_ versus [R10]_total_ (**Figure S4**). As mentioned, the trends are opposite for the two salt conditions. In high salt, [ATP]_coac_ decreases from 3.3 to 0.5 M across [R10]_total_ of 0.5 to 4 mM, while in low salt, [ATP]_coac_ increases but then plateaus to ∼1.3 M starting at 2 mM [R10]_total_ (**Figure 1B**, statistical analysis of comparisons is in **Table S2, S3**).

These findings indicate that, as [R10]_total_ increases, the coacervate phase maintains a more constant local microenvironment for ATP under low salt than high salt conditions. For instance, at low salt, [ATP]_coac_ changes only 2.8-fold, while at high salt [ATP]_coac_ changes 5.7-fold. The [R10]_coac_ is well buffered at both salt concentrations. To gain further insights into thermodynamics of R10 and ATP partitioning via phase separation, we next consider changes in partitioning coefficient and Gibbs free energy of R10 and ATP.

### Thermodynamics of coacervate component partitioning

The chemical composition of the coacervate and dilute phases can be used to calculate R10 and ATP partitioning thermodynamic values. Using the data from **Figure 1**, we estimated values for the partition coefficients (Ks) for R10 and for ATP, which is defined as the ratio for concentration for that polyion in the coacervate phase over the dilute phase; i.e. defined for partitioning *into* the coacervate (**Figure S4-5**). The partitioning coefficient for R10, K_P,R10_, decreased with increasing [R10]_total_ for both salt conditions (**Figure S5)**, in both cases corresponding to an increase in [R10]_dilute_ while [R10]_coac_ remained well buffered (**Figure 1A**). The partitioning coefficient for ATP, K_P, ATP_ , decreased with [R10]_total_ at high salt (**Figure S4A**), here due to a stronger decrease of [ATP]_coac_ than [ATP]_dilute_ (**Figure 1B**), but increased with [R10]_total_ at low salt (**Figure S4**), here largely due to a strong decrease in [ATP]_dilute_ with [R10]_total_, with contribution from an increase in [ATP]_coac_ (**Figure 1B**).

The corresponding partitioning Gibbs free energies of R10 and ATP under each experimental condition are provided in **Figure 3**. As expected, values for ΔG_P,R10_ in low and high salt are similar and highly negative, indicating strong partitioning of R10 in coacervates that weakens with increasing the [R10]_total_. Meanwhile, ΔG_P,ATP_ values with respect [R10]_total_ increase in high salt conditions (i.e. weaker partitioning of ATP) but decrease in low salt (i.e. stronger partitioning of ATP). The observed stronger ATP partitioning with more added R10 in low salt implies stronger ionic pairing between ATP and R10 at low salt, enabling stronger ATP partitioning. Compared to this, for high salt condition, increasing R10 is unfavorable for ATP enrichment in the coacervate phase.

**Figure 3.**
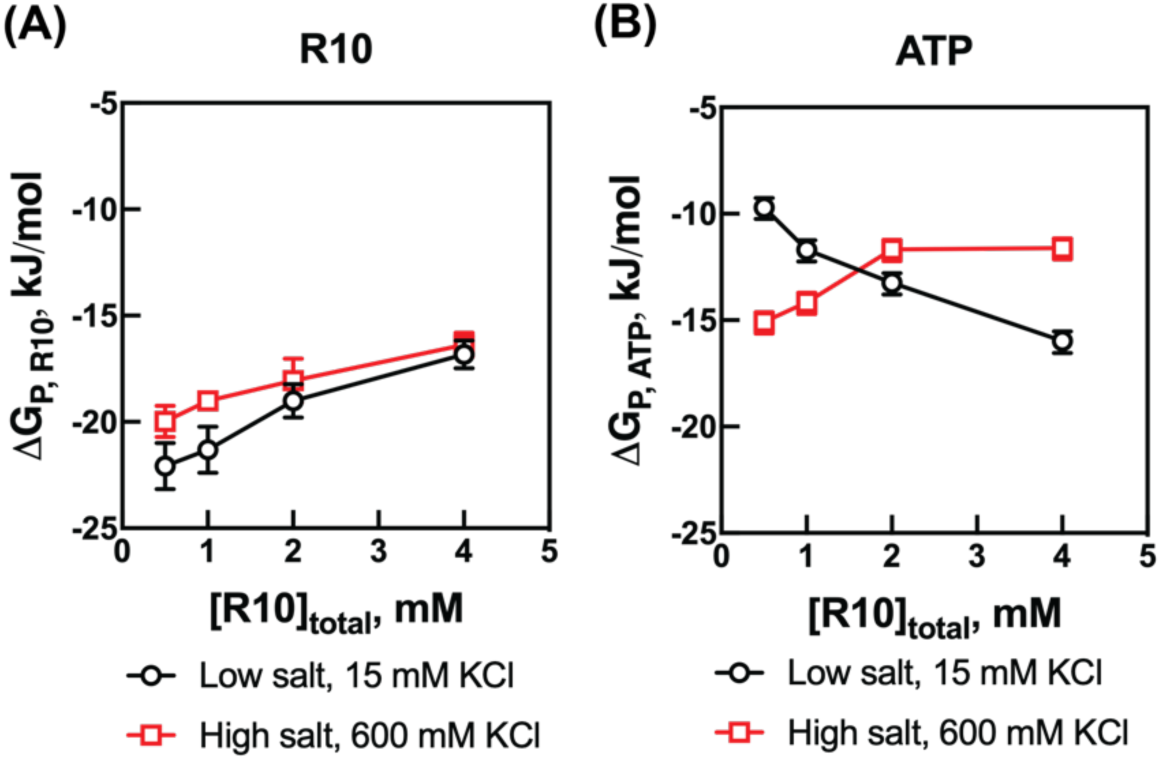
Gibbs free energy of R10 and ATP partitioning in coacervates. Gibbs free energy of partitioning of (**A**) R10 (ΔG_P,R10_) and (**B**) ATP (ΔG_P,ATP_) as a function of [R10]_total_. ATP concentrations are fixed at 10 mM (40 mM_c_ in charge at pH 8.5). Data points are calculated by using the average values, and error bars are the error propagation values from standard deviation in Figure 1. Free energy values are provided in **Table S1, S2.** The y-max and y-min are the same on both panels to facilitate comparison between partitioning free energies of different polyions.

We interpret this crossover in ATP partitioning thermodynamics to be a result of shifting the relative importance of the intermolecular interaction modes between coacervate forming molecules. At high salt, the ion pairing interactions between oppositely-charged molecules (R10–ATP) is less favored due to lower entropy gain by releasing counterions into a high counterion background (i.e. less entropy gain from mixing) as compared to low salt condition. On the other hand, other interaction modes, particularly *π*-*π* interactions between ATP molecules ^51, 52^ and Arg-Arg side chains, and potentially cation-*π* interactions between Arg-ATP, become stronger at higher solution ionic strength^35^. At low salt, repulsions between like-charged molecules (R10–R10, ATP–ATP) is likely hindering other interaction modes. Changes in the relative amounts of R10 and ATP in the coacervate phase can be expected to alter the compartment’s microenvironment, potentially impacting incorporation and function of solutes such as RNA.

### RNA compartmentalization

For coacervates to serve as functional RNA compartments, they need to localize RNA, a phenomenon that is well-known in extant biology^53^. The increased coacervate volume fraction seen with increasing [R10]_total_ (**Figure 2, S3**) is expected to impact local concentrations of guest molecules such as RNAs. If all else remains unchanged –specifically, [RNA]_total_, total volume, and RNA partitioning coefficient (K_RNA_ = [RNA]_coac_/[RNA]_dilute_)– then increased coacervate volume fraction should result in reduced [RNA]_coac_.^25, 54, 55^ Additionally, the observed changes in [ATP]_coac_ (**Figure 1, 3**), could impact RNA partitioning. We thus sought to understand how RNA partitioning changed as a function of increasing [R10]_total_ in the low and high salt R10/ATP systems.

We evaluated both double-stranded and single-stranded RNA oligonucleotides (dsRNA and ssRNA, respectively), which mimic folded and unfolded RNAs, respectively. We used a 10mer RNA that does not fold, 5′-ACCUUGUUCC[Alexa488]-3′, RNA-Alexa488, and its complementary strand (see Methods for RNA design strategy, **Table S4**). We chose to study both dsRNA and ssRNA for several reasons. While dsRNA has higher charge density, suggesting it could exhibit stronger ion pairing interactions with R10 –particularly at low salt, its longer persistence length may physically limit entry to some coacervates and the distance between the bases in A-form dsRNA is too small to allow intercalation of an amino acid^56^. On the other hand, ssRNA is more flexible and its nucleobases more accessible for cation-*ρε*, *ρε*-*ρε*, and/or hydrogen bonding with the guanidinium groups of R10.^57^ Considering the impact of each condition on R10-ATP binding can be useful here, as RNA must compete for binding sites on R10 with the 10^5^ times more abundant ATP molecules (0.1 µM RNA and 10 mM ATP in all samples).

**Figure 4** shows RNA accumulation in coacervates as a function of R10/ATP charge ratio at the two salt conditions. We added 0.1 μM RNA to the total volume. In all cases, RNA partitioned strongly into the coacervate droplets, with estimated local concentrations ([RNA]_coac_) ranging from ∼5 to 55 μM inside the coacervates, an enrichment of 50- to 550-fold. As [R10]_total_ increased, [RNA]_coac_ decreased for both ssRNA and dsRNA and both salt conditions, which is broadly consistent with the increase in relative volume fraction of the coacervate phase noted above (**Figure 2, S3**). Despite these similarities in behavior at low and high salt concentrations, we did observe differences. In the low salt condition, dsRNA reached somewhat higher concentrations in the coacervates as compared to ssRNA, consistent with the higher charge density of dsRNA and with prior work highlighting the importance of ion pairing interactions in RNA partitioning ^18, 58^.

**Figure 4.**
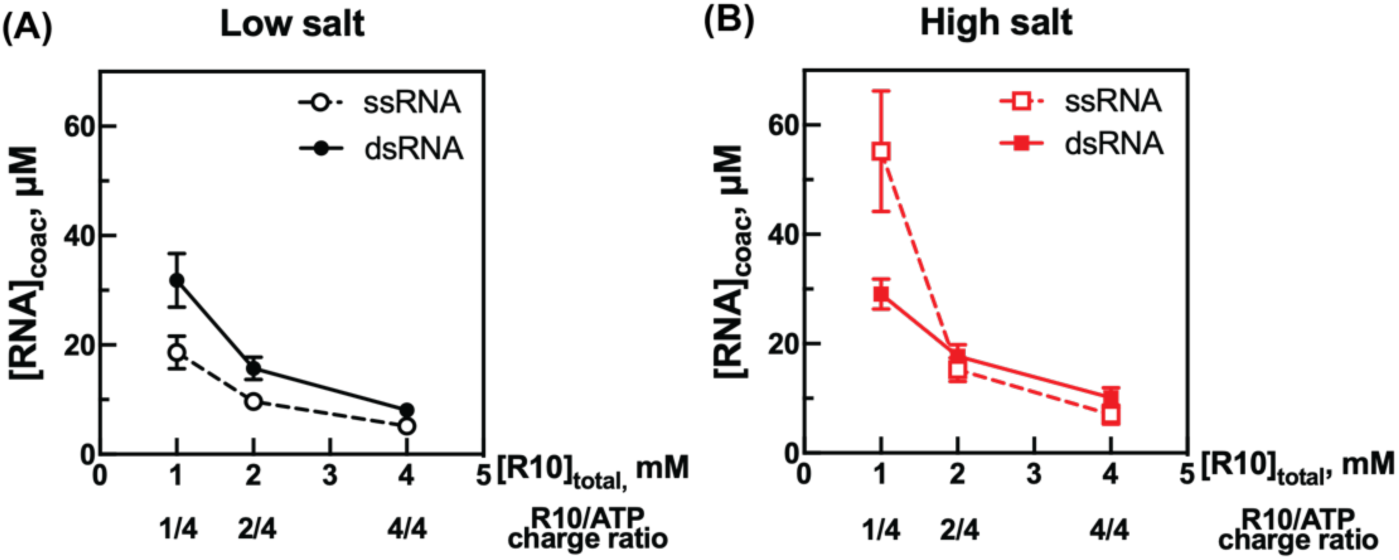
RNA accumulation in R10:ATP coacervates. Quantification of ssRNA and dsRNA concentrations in coacervates as a function of [R10]_total_ at a fixed ATP concentrations (10 mM or 40 mM_c_ in charge at pH 8.5) under **(A)** low salt condition (15 mM KCl, 0.5 mM MgCl_2_) and **(B)** high salt condition (600 mM KCl, 0.5 mM MgCl_2_) with 0.5 mM MgCl_2_ and pH 8.5. ssRNA-Alexa488 was mixed with coacervate samples to be 0.1 µM in total. Local RNA concentration inside coacervates was estimated based on ssRNA-Alexa488 as described in Methods and in **Figure S6.** The average values and standard deviation of RNA concentration are from total 45 ROIs of three independent samples (**Table S5**). Statistical analysis results of pairwise t-test, Pearson’s correlation test and Spearman’s rank test are stated in **Table S6-S7.** For panels A and B, the y-max and y-min are the same on both panels to facilitate comparison of RNA partitioning between salt conditions.

Turning to the high salt condition, as mentioned we find that both [ssRNA]_coac_ and [dsRNA]_coac_ again decrease with increasing [R10]_total_, consistent with the larger coacervate volumes as [R10]_total_ increases. Counterintuitively, despite the importance of charge-charge interactions between R10 and RNA for partitioning, local RNA concentrations in the coacervate phase were not reduced at high salt as compared to low salt, with [dsRNA]_coac_ nearly the same and [ssRNA]_coac_ higher for the high salt condition. This observation can be rationalized by higher salt reducing charge-charge interactions, including those between R10 and ATP, such that RNA is able to displace the ATP molecules from the coacervate to interact with R10 molecules. Finally, at high salt we observed significantly higher [ssRNA]_coac_ than [dsRNA]_coac_ at only the lowest [R10]_total_. Higher partitioning for ssRNA than dsRNA is consistent with greater accessibility of the nucleobases in unpaired RNA allowing them to interact with R10, which is particularly important at high salt where the ion pairing interactions are weaker. Why then is [ssRNA]_coac_ not also higher than [dsRNA]_coac_ for the 2/4 and 4/4 charge ratios? We interpret this as resulting from the change in composition and properties of the coacervate phase, specifically the reduced [ATP]_coac_ and increased R10-R10 interactions, in addition to the aforementioned volume and ionic strength effects.

Overall these observations for RNA accumulation as a function of R10:ATP ratio are consistent with an interpretation that at low salt, where ion pairing interactions are important both in coacervation itself and in the accumulation of “guest” RNA molecules, changes in coacervate volume allow the chemical microenvironment of the coacervate phase to be largely maintained. Guest molecule concentrations however are impacted by the change in relative phase volumes, with larger coacervate volumes decreasing local RNA concentrations. In contrast, for the high salt condition, additional intermolecular interaction modes become important and the composition and properties of the coacervate microenvironment also change with changing availability of coacervate-forming molecules. Consequently, the impact on guest molecule uptake is less straightforward.

### Effect of pH on RNA partitioning

To further test RNA partitioning by the balance of coacervate phase volume effect and altered intermolecular interactions, we challenged the system by lowering pH. In this way, we can also evaluate scenarios plausible in hydrothermal spring water or acidic oceans. For example, the above experiments were performed at pH 8.2 ∼ pH 8.5 ± 0.1, which is within the range of pH values for spring water and ocean water.^22, 59–61^ Since lower pH waters could also be important for prebiotic chemistries, we also evaluated RNA accumulation in R10/ATP coacervates at pH 6.5 ± 0.2 and 4.5 ± 0.2, which can represent hydrothermal spring water or acidic oceans.^22, 60, 61^ Across this pH range, ATP is expected to change its singly protonated state, gaining a proton on the gamma phosphate partially at pH 6.5 and fully at pH 4.5 and partially on its nucleobase atom N1 at the lowest pH (**Figure S7**)^62, 63^. Within the tested pH range in this study, the net charge of ATP changes from –4 at ∼ pH 8.5, – 3.5 at pH 6.5) to – 2 at pH 4.5.

We anticipated that decreasing the net charge of ATP by 1 to 2 units at the lower pH values would weaken its interactions with R10 and thus allow RNAs to compete better for interactions with R10, which could result in stronger RNA partitioning. We note that our RNAs lack a terminal phosphate (pKa 6.5); all other phosphates are negatively charged within the entire tested pH range (phosphodiesters pKa of 1), giving them an advantage over ATP for ion pairing with R10 as pH is lowered. Additionally, at pH 4.5, adenine and cytosine nucleobases (both have pK_a_ on the imino nitrogens of ∼4.2)^62^ could begin to protonate in ssRNA, although not dsRNA, where base pairing shifts p*K*_a_s further from neutrality.^64^

All tested charge ratio, ionic strength, and pH conditions led to coacervation, making it possible to compare RNA localization across the full set of solution environments (**Figure 5A, B,** microscope images of 2/4 R10/ATP charge ratio are in **Figure S9**). A markedly different pH response of RNA partitioning was observed depending on the solution ionic strength. At low salt, the [RNA]_coac_ was insensitive to pH, while at high salt, [RNA]_coac_ increased sharply at low pH (**Figure 5**, statistical analysis for pairwise t-tests, Pearson’s correlation test, Spearman’s rank test are in **Table S8, S9**). The same general trends were observed for ssRNA and dsRNA (**Figure 5C, D**). We again consider the impact of each condition on R10-ATP binding as above, as RNA must compete for binding sites on R10 with the more abundant ATP molecules. Due to the lower multivalency of ATP as compared to either ss or dsRNA, increased ionic strength can be expected to disrupt R10 binding to ATP more so than to RNA, which could result in stronger [RNA]_coac_ under high salt conditions as observed (**Figure 5B**). As shown by recent theoretical calculations,^65^ a loss of polyanion charges with decreasing pH, which can occur here at pH 4.5 from protonation of gamma phosphate and adenosine of ATP, could be compensated by increased amount of polyanion and free anion in the coacervate phase of a polyelectrolyte complex coacervate system to maintain net charge neutrality in coacervates. In our system, charge neutrality in the coacervate phase could be maintained at lower pH by incorporating additional ATP molecules, RNA, and/or Cl^-^ into the coacervate.

**Figure 5.**
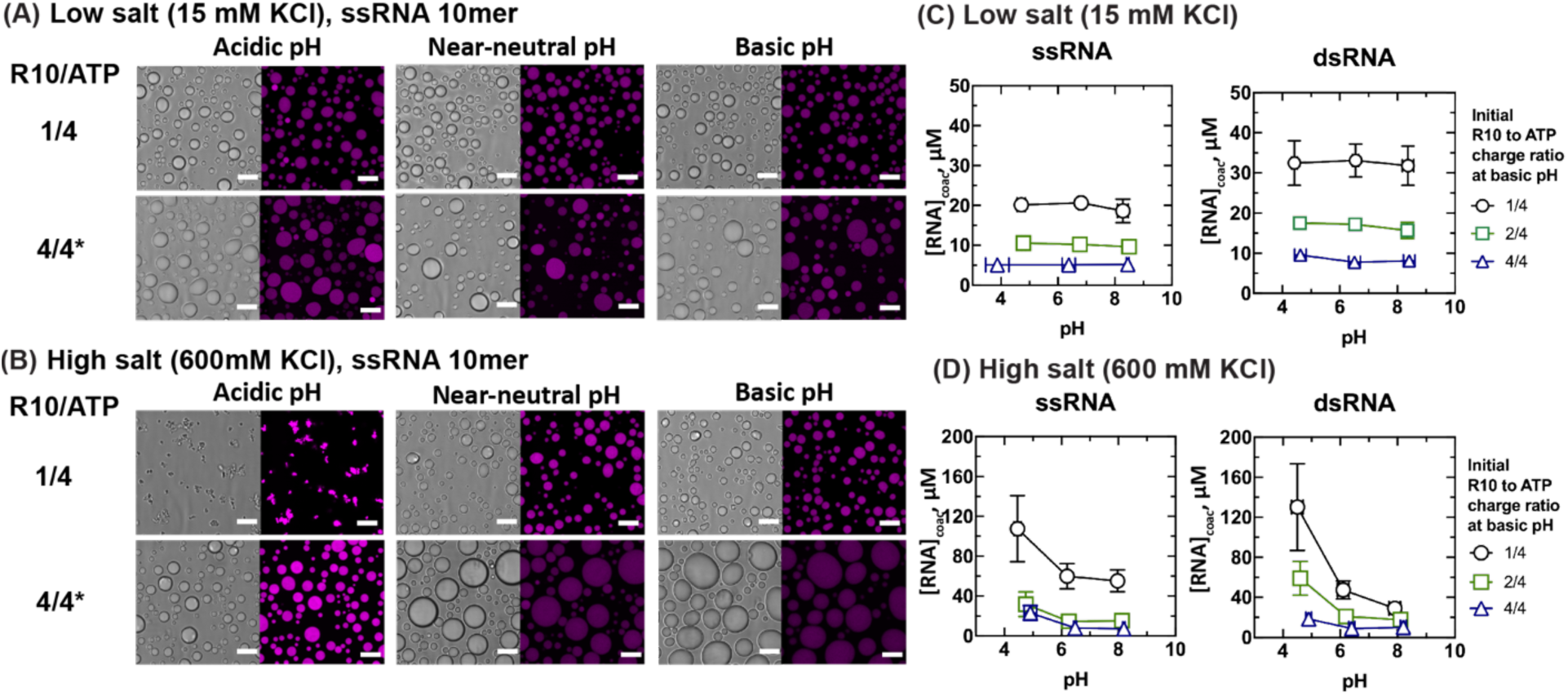
Fluorescent RNA partitioning in coacervates is sensitive to pH at high salt. Confocal microscope images of ssRNA-Alexa488 of 1/4 and 4/4 R10/ATP initial mixing charge ratio (determined at basic pH) in **(A)** low salt and **(B)** high salt conditions. Grey-colored images are by bright-field microscopy, and magenta-colored images indicate the fluorescence intensities from the confocal microscopy. “4/4*” indicates that intensity of microscope images of 4/4 R10/ATP images were enhanced in same degree within the same salt condition to aid visualization. Raw microscope images are provided in **Figure S8.** ssRNA-Alexa488 was mixed with coacervate samples to be 0.1 µM in total. Quantification of ssRNA or dsRNA concentrations in coacervates of various R10 to ATP charge ratios in **(C)** low salt condition (15 mM KCl, 0.5 mM MgCl_2_) or **(D)** in high salt condition (600 mM KCl, 0.5 mM MgCl_2_). Intensity values from confocal images are converted to RNA concentration using the calibration curves in **Figure S5**. Scale bars indicate 10 µm. The average values and standard deviation of RNA concentration are from total 45 ROIs of three independent samples (**Table S5**). Statistical analysis results for pairwise comparisons, Pearson’s correlation test, and Spearman’s rank test are stated in **Table S8-S9.**

### Fraction of total RNA encapsulated in coacervates

Notably, both salt conditions show higher local RNA concentration in coacervate phase under limiting [R10]_total_ (**Figure 4**), which happens despite the insensitivity of [R10]_coac_ to [R10]_total_ seen in **Figure 1A**. Indeed, [RNA]_coac_ values are roughly 4x and 2x greater for 1/4 R10/ATP or 2/4 R10/ATP charge ratios, as compared to [RNA]_coac_ in 4/4 R10/ATP of similar pHs (**Figure 4**). As described in the previous sections, this can be understood in light of how the volume fraction of coacervate phase impacts [RNA]_coac_ at a constant partitioning coefficient.^25, 55^ As more R10 is added to excess ATP, the volume of the coacervate increases in a manner proportional to [R10] (**Figure S3**). This has the effect of diluting the [RNA]_coac_ for 2/4 and 4/4 as compared to 1/4 R10/ATP. Theoretically, the local [RNA]_coac_ is inversely proportional to the coacervate phase volume at constant partitioning coefficient K_RNA_, [RNA]_total_, total sample volume, and RNA encapsulation efficiency (*f*_coac_= N_coac_/N_total_, where N_coac_ is moles of RNA in the coacervate phase and N_total_ is total moles of RNA in sample).^25, 31^ In such cases, the fraction of total RNA molecules that are sequestered within the coacervate phase could be more indicative of such environmental changes than their local concentrations.

To determine the encapsulation efficiency of the RNA (*f*_coac_), we 5′-end ^32^P-radiolabeled the RNA and then determined the RNA counts in the coacervate and dilute phases and estimated *f*_coac_ (see Methods). Liquid scintillation counting of radiolabeled-RNA provides excellent dynamic range and sensitivity and can accurately determine number of RNAs in dilute phase without the influence of fluorescent labels on partitioning. **Figure 6** summarizes the *f*_coac_ values in the different charge ratios and salt and pH conditions from **Figure 5**. Under all tested conditions, most of the total RNA was localized within the coacervate phase, despite its small relative volume (<1% of total volume, see **Figure S3**). Measured *f*_coac_ values ranged from as high as ∼0.98 for dsRNA under low salt, either pH, and 4/4 charge ratio conditions to as low as ∼ 0.70 for ssRNA under high salt, low pH and 1/4 charge ratio conditions.

**Figure 6.**
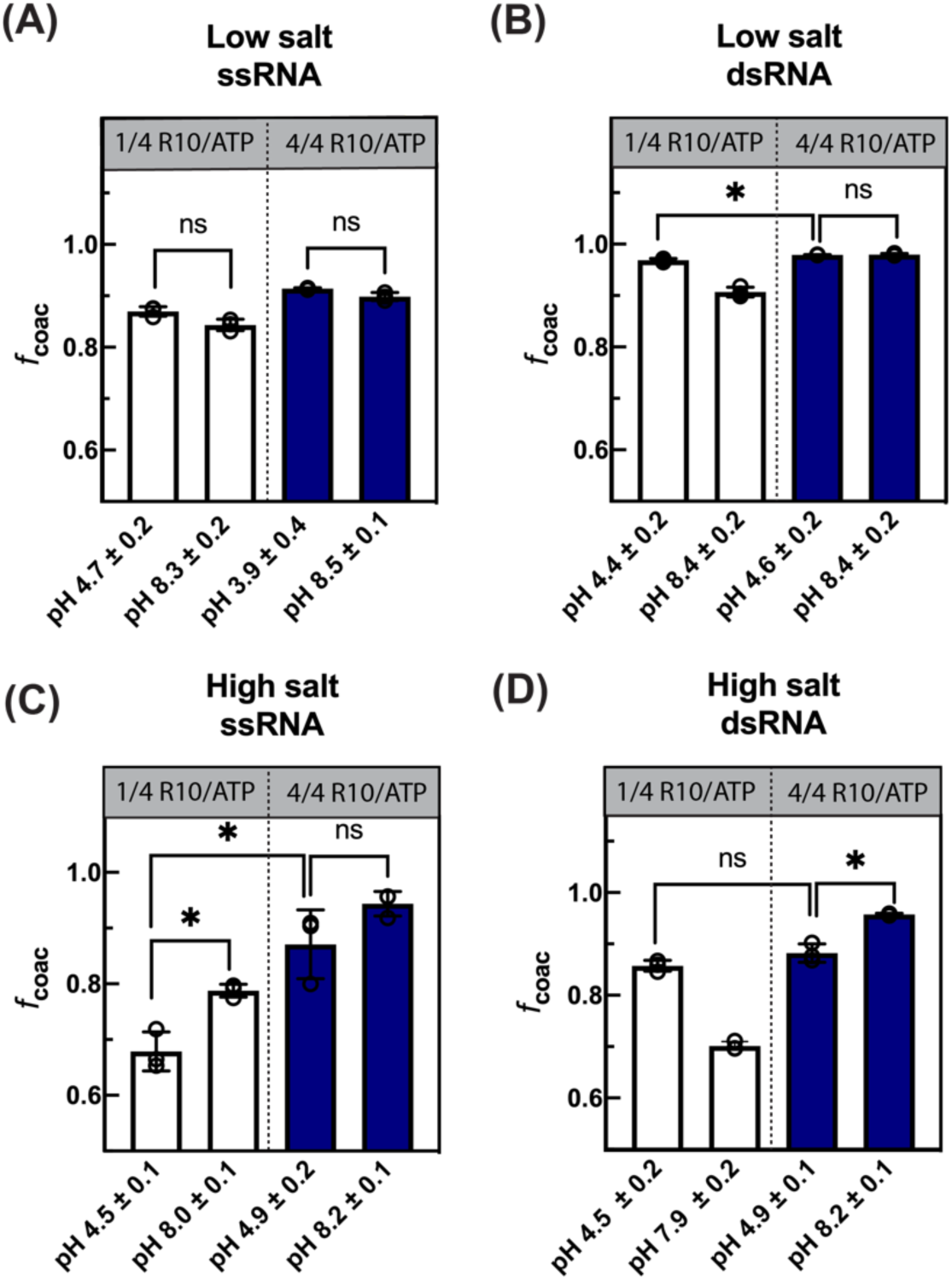
Fraction of RNA within coacervates as a function of salt, pH, and R10/ATP charge ratio from radiolabeling experiments. RNA fraction in coacervates in low salt conditions: **(A)** ssRNA and **(B)** dsRNA. RNA fraction in coacervates in high salt conditions: **(C)** ssRNA and **(D)** dsRNA. Error for pH values indicate the standard deviation of three independent pH measurements by pH meter. Three independent trials of fraction of RNA in coacervates are shown as symbols. Bars indicate the average, and the error bars indicate the standard deviation (**Table S10**). We note pair-wise p values in the graph that are “ns” or “*”, while rest of pair-wise P-values are statistically significant and listed in **Table S11-S14** (they are **, ***, and ****). *: p value < 0.05, **ns**: p value > 0.05 (statistically not significant).

Notably, the strong trends of higher local [RNA]_coac_ for higher salt as compared to lower salt, especially at low pH or at lower R10/ATP charge ratio seen in **Figure 5** are not associated with more RNA strands overall being sequestered in the coacervate phase. Indeed, the *f*_coac_ data show the opposite trends, with *f*_coac_ *decreasing* at high salt and *increasing* with [R10]_total_ (**Figure 6**). As 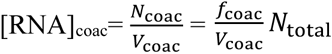, the opposing trends in [RNA]_coac_ and *f*_coac_ can be partially understood as a consequence of differences in the coacervate volumes. As more of the limiting R10 is added to the system, formation of a <u>proportionally increased</u> coacervate phase (e.g. increase in V_coac_) (**Figure S3-4**), along with a <u>decreased</u> local concentration of coacervate-sequestered RNA (**Figure 4**), enables even more total RNA strands to be sequestered giving an <u>enhanced increase</u> in *f*_coac_ with [R10]_total_, especially at high salt (**Figure 6**).

Differences in the pH dependence of *f*_coac_ were apparent between the low and high salt samples. RNAs were well sequestered in the coacervate phase regardless of conditions of pH and charge ratio under low salt (*f*_coac_ ranges from ∼ 0.87 to ∼ 0.98, **Figure 6 (A)-(B)**), which seems overall consistent with our observation from RNA-Alexa488 experiments in **Figure 5**. Under high salt conditions, *f*_coac_ for dsRNA in 1/4 R10/ATP coacervates was higher in acidic pH consistent with the observation of much higher local RNA concentration in coacervates seen in **Figure 5**. At 4/4 R10/ATP in high salt, *f*_coac_ values of both dsRNA and ssRNA were lower or similar in acidic pH as compared to basic pH (**Figure 6(C)-(D))**. These observations can again be understood in terms of relative phase volume differences: coacervate phase volumes are larger in 4/4 R10/ATP as compared to 1/4 R10/ATP, and larger in high pH overall (**Figure S3, S11,** again, 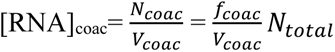). Overall, the complementary [RNA]_coac_ and *f*_coac_ measurements, in Figures 4 and 5, support three important observations: **(1)** Strong RNA compartmentalization was observed under all tested conditions in both salt conditions. **(2)** RNA partitioning levels are influenced by R10/ATP charge ratio in all tested pH/salt. **(3)** RNA accumulation is more sensitive to pH under high salt conditions. We conclude that these observations could stem from the multiple factors affecting RNA distribution in multiphase systems, in particular the balance of interactions between R10-RNA versus R10-ATP, and the impact of fractional coacervate phase volumes.

### RNA duplex stability in R10/ATP coacervates

As RNA partitioning levels are well retained in all tested conditions, to test their potential functionality as RNA compartments, we examined the stability of RNA duplexes to those ranges of conditions using FRET. We utilized pre-hybridized 10mer dsRNA using a 3′-Cy3-labelled RNA and a complementary 5′-Cy5-labelled RNA as described in Methods and **Table S4.** The dsRNA or ssRNA controls in salt buffer without coacervates are included in the figures, while FRET efficiency values of ssRNA in buffer are all close to 0. Under low salt conditions, significantly reduced FRET efficiency of dsRNA was seen for coacervate-compartmentalized RNA at both pH 8.4/8.5 and 4.7 relative to dsRNA buffer controls, indicating some duplex destabilization (**Figure 7A, Table S15).** We attribute the weakened RNA duplex association in these R10/ATP systems as resulting from favorable interactions between RNA nucleobases in the single-stranded states and guanidinium sidechains of R10,^66^ which are present at nearly 10 M (**Figure 1A**). The FRET efficiency was insensitive to pH at low salt but was sensitive to the molecular charge ratio of R10/ATP (**Figure 7A, Table S18** for pairwise p-values). In particular, 4/4 R10/ATP samples showed weaker duplex association for both salt conditions giving 18 % to 27 % lower FRET efficiency than 1/4 R10/ATP coacervates at the same pH/salt condition, consistent with greater availability of R10 sidechains to interact with ssRNA (**Figure 1**) and/or the lower local RNA concentration inside the 4/4 R10/ATP samples (**Figure 5B**).

**Figure 7.**
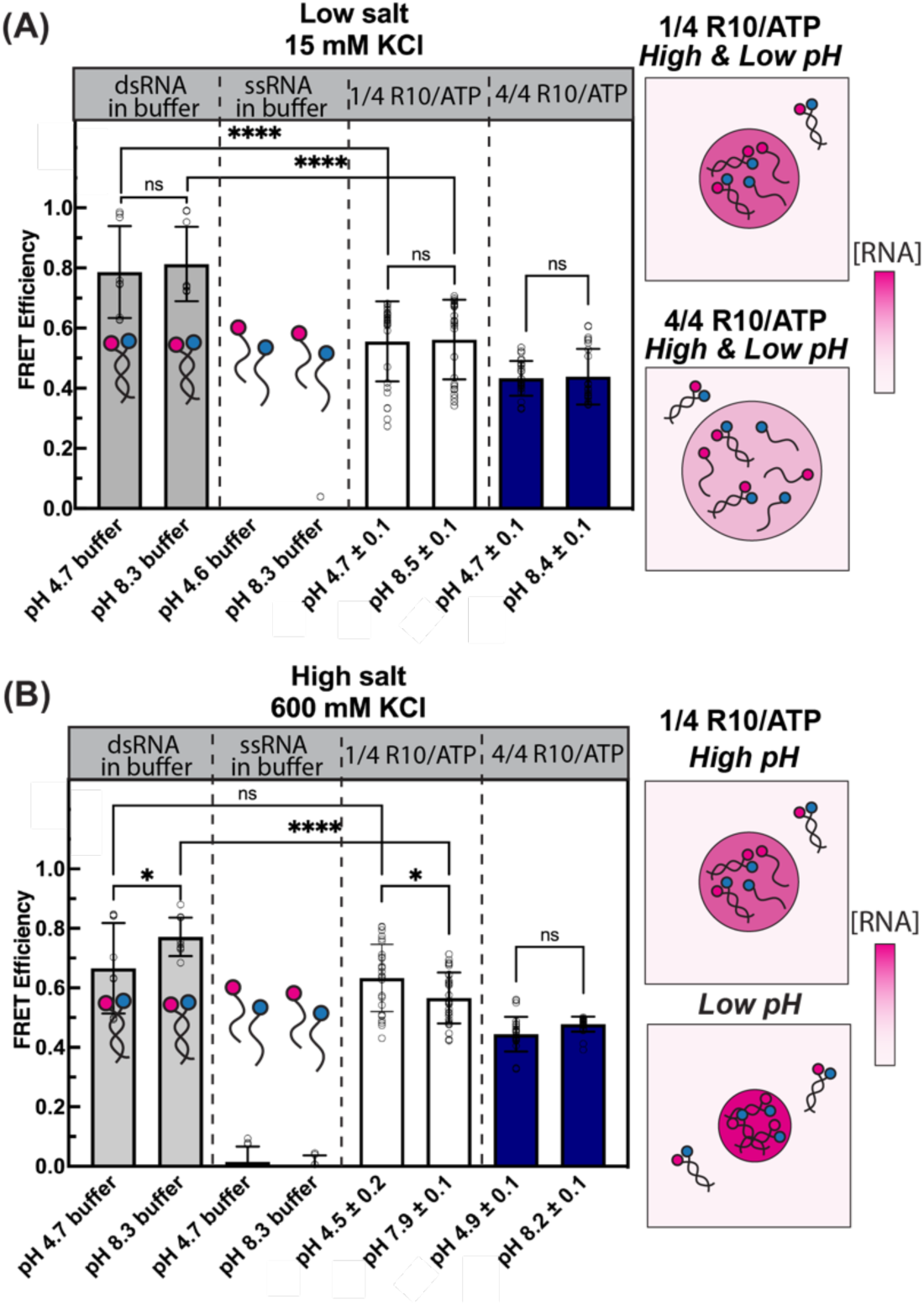
FRET efficiency of RNAs in R10/ATP coacervates. FRET efficiency values at low and high salt conditions are plotted in panels **(A)** and **(B)**, respectively. White bars indicate samples of 1/4 of R10/ATP mixing charge ratio with indicated pHs on x-axis. Blue bars are samples of 4/4 of R10/ATP mixing charge ratio with indicated pHs on x-axis. Bars and error bars indicate the average values and the standard deviations of all ROIs from three independent samples (27 ROIs for coacervate samples and 9 ROIs for dsRNA or ssRNA in salt buffer without coacervates) (**Table S15**). We used *⍺* and *β* values for FRET efficiency calculation as stated in Method and **Table S16-S17**. We denote “ns” or “*” in the graph, while rest of pair-wise P-values are statistically significant and listed in **Table S18-S19.** *: p value < 0.05, **ns**: p value > 0.05 (statistically not significant).

Patterns of RNA duplex stabilities under high salt conditions were similar to those under low salt conditions. An exception could be the 1/4 R10/ATP coacervates in high salt, which showed slightly higher FRET efficiency for the low pH condition and was statistically indistinguishable from buffer at the same pH (**Figure 7B, Table S19** for pairwise p-values). This can be understood in terms of the higher local RNA concentration in the low pH, high salt 1/4 R10/ATP coacervates (**Figure 5D**) and stronger *π*-*π* interactions, which could help stabilize against dissociation despite strong interactions between the RNA and the R10. Overall, our results indicate that RNA dissociation thermodynamics in R10/ATP droplets are largely maintained under pH and salt changes, while being more sensitive to changes in R10/ATP ratio.

## Conclusion

Phase systems based on multiple molecules and/or multiple interaction types are only just beginning to be studied in relation to molecular buffering. We examined coacervates formed from the relatively low molecular weight components, R10 and ATP, under prebiotically inspired salt and pH conditions, as well as different mixing ratios of R10 and ATP. These environmental variables affect interactions between coacervate molecules themselves and their interactions with guest molecules like RNAs. We found that, while some observations can be understood in terms of relatively straightforward increases to the relative volume of coacervate phase as [R10]_total_ is increased, others cannot. Molecular buffering of R10 was effective under both salt conditions, while local ATP concentrations were somewhat more variable. A relatively better molecular buffering of local ATP and RNA concentrations were achieved under low salt conditions. Although ATP and RNA buffering was less effective at high salt, this condition led to stronger enrichment of ATP and RNA in the coacervates. This can be understood in terms of how KCl concentration influences the dominant intermolecular interaction modes between coacervate-forming molecules and between these molecules and guest RNAs. As KCl concentration increases, ion pairing interactions between R10 and ATP become less important, while allowing ρε-ρε interactions to become more important, making self-interactions less unfavorable (R10-R10 and ATP-ATP).^35^ This, combined with the coacervate phase volume effect, could aid in localizing ATP and RNA even under limiting [R10]_total_.

The system can maintain molecular buffering of R10 even in high salt condition, although ATP buffering is not achieved. This can be understood in terms of shifting contributions of self- and non-self-associating interaction modes, more specifically by considering how changing ionic strength alters the balance of repulsive and attractive R10–R10 (self) interactions and the relative strengths of attractive R10-ATP (non-self) interactions. Theoretical investigations have suggested that altering interaction modes of phase separating molecules could influence the efficiency of concentration buffering.^29^ Buffering efficiency is shown to increase when protein condensates are formed only by self-associating (homotypic) interactions^27^, while protein biomolecular condensates formed by multiple components via non-self-associating (heterotypic) interactions do not exhibit a fixed buffered concentration^30^. The relationship between molecular interactions and molecular buffering efficiency for biological compartments remains unclear, but weakening heterotypic interactions could help maintain the buffering ability of phase separating systems either in coacervate phase or in dilute phase under molecular concentration perturbations.^27, 29, 30^ This can in part support our observation that salt weakening R10-ATP interactions can lead to sustained R10 buffering while worsening the ATP buffering (as ATP begins to act less like a coacervate component and more like a guest molecule). Whether one scenario (e.g., holding [R10]_coac_ constant) is more advantageous than another (e.g., maintaining constant R10/ATP ratio in the coacervate, or maximizing RNA compartmentalization, etc) depends on context. With growing interests in various interaction modes in proteins with nucleotides for phase separation,^35, 44, 67^ our study on simple model systems of oligoarginine and ATP can contribute to elucidate the consequences of such interaction modes in phase separation^27, 30^.

Our findings in these simple oligoarginine and ATP systems suggest mechanisms by which coacervates modulate their roles as buffering compartments under different environmental conditions (e.g., ionic strength, pH). In general, changes in coacervate phase volume can accommodate variations in availability of phase-separating molecules while largely maintaining the same coacervate microenvironment, a process that inherently alters the local concentration but not necessarily overall abundance of guest molecules (in our system, RNAs) within the coacervates. Here, as [R10]_total_ increases, local [RNA]_coac_ decreases, even as the total amount of compartmentalized RNA increases. Enticingly, by shifting interaction modes in response to solution condition changes, coacervates can enrich local RNA concentration further. This can lead to different types of primitive molecular buffering depending on whether the compartments are located in lower salt conditions (groundwater and hydrothermal springs) or in higher salt conditions (seawater and hydrothermal vents). For example, a more consistent local RNA concentration could be maintained within the compartments despite changing pH in spring water or lakes, while compartments with stronger localization of RNA could be generated in weakly acidic salty water. This interplay between relative strengths of dominant intermolecular interaction modes and coacervate phase volume changes determine its functionality as compartments that can adapt to molecular concentration fluctuations and/or enrich guest molecules. When the steady state of living cellular systems is disturbed by external factors, it can actively maintain its homeostasis via regulatory mechanisms, which is a guiding principle of all extant life.^68^ The ability of even low multivalency coacervates to exhibit molecular buffering is an important step towards a primitive homeostasis that could have been important in emergence of life.

## Acknowledgments

This work was supported by the NASA Exobiology program grant 80NSSC17K0034 and no. 80NSSC22K0553 (S.C., M. O.M., P.C.B., and C.D.K.). S.C. was also supported by Future Investigators in NASA Earth and Space Science and Technology (FINESST) under Grant 80NSSC19K1531.

## Additional Information

Supplementary Information is available in the online version of the paper.

## Author Contributions

M. O. M. performed the radiolabeled RNA partitioning experiments. S. P. L. performed the ATP and R10 quantifications of high salt conditions and single stranded RNA partitioning experiments. S.C. performed all other experiments. All authors conceived and designed the experiments and analyzed the data. S. C., P.C.B., and C. D. K wrote the manuscript with input from S. P. L. and M. O. M.

## Experimental section

### Materials

Poly(L-Arginine hydrochloride) (degree of polymerization n = 10, 95% purity or above) (R10) was purchased from Alamanda Polymers and used without further purification. Adenosine Triphosphate (ATP) was purchased from MP Biomedicals as disodium salt hydrate form (CAS No. 51963-61-2). N-terminus TAMRA labeled version of R10 was custom-synthesized from Biomatik. RNA and RNA-Alexa488 are purchased from IDT. All other oligoRNAs for FRET analysis are custom-synthesized and purchased from Sigma Aldrich.

### Preparation of coacervate samples

R10 stock solutions were prepared as 10 mM final concentration in HPLC grade water (final pH is around 8.3). ATP stock solution was prepared as 100 mM final concentration and pH was adjusted to pH 8.3 using 1 M NaOH. Aliquots of peptide and ATP stock solutions were sealed with inert gas in and kept under -22 °C for storage. Before use, stock solutions were thawed and vortexed thoroughly. Coacervate samples are always freshly prepared before characterizations. We prepared samples with the following order of addition: HPLC water, salt buffers, ATP stock solution, fluorescent-labeled RNA or fluorescent-labeled peptide, and R10 (None of fluorescent molecules are added for ATP quantification experiments). Coacervate samples were equilibrated at room temperature for 15 mins before analysis. ATP concentration was fixed to 10 mM (40 mM in charge concentration estimating as– 4 charges in total at pH 8.3), while R10 concentration was varied from 4 mM to 0.5 mM (40 mM to 5 mM in charge concentration estimating as +10 charges in total at pH 8.3). Desired amounts of R10 and ATP were mixed in either one of two different salt buffers: low salt buffer (15 mM KCl and 0.5 mM MgCl_2_) or high salt buffer (600 mM KCl and 0.5 mM MgCl_2_). We fixed the magnesium concentration to 0.5 mM MgCl_2_ to focus on the effect of salinity not the divalent cation effect in our experiments. The pH of the coacervate samples is measured by Mettler Toledo Ultra Micro ISM electrode after at least 1 hour of equilibration followed by 30 mins of centrifugation, as turbid coacervate samples sometimes give large fluctuations in pH readings.

### Microscopy and image analysis

Confocal images were taken using a Leica TCS SP5 laser scanning confocal inverted microscope (LSCM) with Leica LAS AF software and an HCX PL APO CS 63.0×/1.40 oil UV objective. Fluorescence intensity values were acquired from raw fluorescence images using Fiji^69^ after background correction.

### R10 partitioning by confocal microscope or fluorimeter

TAMRA-labeled R10 were mixed with non-fluorescently labeled peptides to be 0.0125 ∼ 0.1% TAMRA-labeled strand by concentration (fixed as 0.5 uM TAMRA-R10 total). Low percentage of fluorescently labeled R10 was used to minimize artifacts.^70, 71^ Coacervate phase R10 concentration were estimated from intensity of TAMRA-R10 measured by confocal microscope using 543 nm laser and collected from 560 – 660 nm with a DD 488/543 dichroic filter. Dilute phase R10 concentrations were estimated by measuring TAMRA-R10 in dilute phase by fluorimeter; emission of TAMRA-R10 from 560 nm to 700 nm was collected by a Jobin Yvon Horiba FL3-21 fluorimeter with 5 nm slit size, 5 average scan, and 543 nm excitation. From the confocal microscopy images, intensity of labeled R10 was averaged from 45 droplets total from three independent trials after the background correction. Calibration curves of fluorescent labeled R10 by confocal microscope and fluorimeters were acquired with the same confocal microscope or fluorimeter setting as a function of fluorescent labeled R10 concentrations over fluorescence intensity (**Supplementary Figure 1**). Stock solutions of fluorescent labeled R10 in water were used for the calibration by diluting it with water because calibration could not be performed directly in the coacervate media. TAMRA should be relatively insensitive to differences between buffer and coacervate media but some impact on emission intensity cannot be ruled out^72^. Using the calibration curves, we calculated the concentration of fluorescent labeled R10, and then, used the dilution factor to estimate the concentration of charged groups from R10.

### Coacervate phase volume estimation

We found that different wetting properties of solutions of oligoarginine and ATP in various pH and salt conditions hinder the accurate volume estimation. Therefore, we estimated the coacervate phase volume by concentration of R10 in dilute phase and coacervate phase as shown in Figure 1 using the following equation (Supplementary Figure 2): 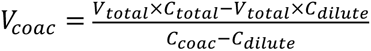, where *V_coac_*is coacervate phase volume, *V_total_* = total volume, *C_total_* is total R10 concentration in solution and C_dilute_ is R10 concentration in dilute phase. We confirmed the trend of coacervate volume changes from this estimation to that of comparing sizes of coacervate phase volume to the known reference volumes (Supplementary Figure 3). Difference of coacervate phase volume percentages by these two estimations is similar for high salt condition but 50 % different for low salt. Nonetheless, comparisons of these two different methods confirms the trends of coacervate phase volume changes as a function of R10 amount in solution regardless of its value difference, allowing comparison of coacervate phase volumes of charge ratios of R10/ATP.

### ATP partitioning by UV-vis spectrometer

ATP concentration in dilute phase were estimated from 260 nm absorbance using the Beers-Lambert law (1 cm quartz cuvettes, ATP extinction coefficient at 260 nm = 15400 M^−1^ cm^−1^). Dilute phase was removed from coacervate samples after 1 hour equilibration followed by 30 mins centrifugation and then further diluted by HPLC water (200x dilution). ATP concentration in coacervate phase is estimated by ATP concentration in dilute phase and coacervate volume estimated from R10 concentrations from the previous section above using the following equation:

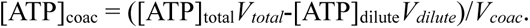

### Partitioning thermodynamics calculation

The equilibrium constant, *K*_i_ is defined as the ratio of the concentration of the molecules of interest (i) in the coacervate and dilute phases: 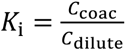, where *C*_coac_ and *C*_dilute_ are the concentrations of those molecules in the coacervate and in dilute phases, respectively. The partitioning Gibbs free energies of R10 (ΔG_p,R10_) and ATP (ΔG_P,ATP_) are estimated based on the equation of Δ*G* = −RT*lnK*, where R is the gas constant (8.314 J mol^−1^ K^−1^) and T is the temperature (here, 298 K).

### Fluorescent-RNA partitioning in coacervates

3′-Alexa488-ssRNA 10mer (5′-ACCUUGUUCC[Alexa488]-3′) are used to study the RNA partitioning in coacervate phases by confocal microscopy. We designed the RNA sequence to have 50 % GC content when double stranded and to avoid self-complementarity by designing one strand (the sense strand) to be pyrimidine-rich and the other strand to be purine-rich (all RNA sequences are listed in **Supplementary Figure 3**). Final total concentration of RNA was 0.1 µM for all samples. For pre-hybridized dsRNAs stock solutions, 3′-Alexa488-ssRNA 10mer and its complementary sequence (5′-ACCUUGUUCC-3′) were mixed to be 0.1 µM each in 150 mM KCl and heated up in 90 °C for 2 mins with following equilibration at room temperature for 1 hour to allow hybridization before they were mixed into coacervate samples. RNA sequences are designed to have 50 % of GC contents and avoid the self-complementary structures as noted in our previous studies.^18^

Calibration curves as a function of concentration of 3′-Alexa488-labeled RNAs over fluorescent intensity were individually achieved with the same setting of confocal microscopes using various concentration of ssRNA-Alexa488 10mer, pre-hybridized dsRNA 10mer from 40 µM to 1 µM. Note that calibration could not be performed directly in the coacervate media; Alexa488 is expected to be relatively unaffected^73–75^ by differences between buffer and coacervate environments, though some impacts on emission intensity cannot be entirely excluded. To minimize errors by mixing, we prepared the mixture of RNA with R10/ATP coacervates in basic pH (∼ pH 8.3) of 150 μl, and then, aliquoted to the 50 μl each, and added different amount of 1 M HCl into each aliquot for desired amount (5 mM HCl or 12.5 mM HCl as a final concentration for ∼ pH 6.5 or ∼ pH 4.5, respectively). We imaged these three samples of different pHs using the confocal microscope within 30 mins of sample preparation.

### Radiolabeling of RNA and its partitioning experiment

5′ Radiolabeling of 10mer and 20mer RNAs was done by kinase reactions prepared with the following conditions: 7 µM of RNA (either 10mer or 20mer strand A), 10 U of T4 Polynucleotide Kinase, 1X PNK buffer and 2.5 µM γ-^32^P ATP. The kinase reaction was incubated at 37 °C for 1 hr before being quenched with 2X formamide loading dye (95% formamide, 20 mM EDTA, 0.1% xylene cyanol, 0.025% bromophenol blue), loaded onto a denaturing 20% acrylamide, 8.3 M urea gel, and fractionated at 20 W for 1 h. The gel was imaged using X-ray film, aligned by using luminescent stickers (Glogos II Autorad Markers from Agilent Technologies), and subsequently the RNA band was excised from the gel, placed in TEN 250 buffer (10 mM Tris pH 7.5, 1 mM EDTA, 250 mM NaCl), and crush and soaked overnight at 4 °C. The next day, it was ethanol precipitated and scintillation counted to determine the concentration of radiolabeled RNA. Radioactive partitioning experiments were performed similarly to those in Frankel et al^32^ and in Choi et al. For ssRNA partitioning experiments, cold RNA and hot RNA were added to a final concentration of ∼ 0.1 uM and ∼50-180 kcpm of radiation per experiment. In dsRNA partitioning experiments, unlabeled RNA and radioactive RNA were added together with KCl, and MgCl^2^ to make a stock solution with 5 uM RNA, ∼50-180 kcpm of radiation/uL, 150 mM KCl, and 5 mM MgCl_2_ and renatured at 95 °C for 2 min. For ssRNA experiments, coacervates were prepared by addition of the following components: water, low salt buffer (15 mM KCl, 0.5 mM MgCl_2_), ATP, a mix of unlabeled and radioactive RNA, and R10 for a total reaction volume of 50 uL. For dsRNA experiments, coacervates were prepared by addition of the following components: water, low salt buffer (15 mM KCl, 0.5 mM MgCl_2_), ATP, a mix of unlabeled and radioactive RNA, and R10 for a total reaction volume of 50 uL. Once the coacervates were formed, they were allowed to equilibrate for 15 minutes to 1 hr in order to allow the partitioning of the RNA to occur. After equilibration, they were centrifuged at 17,000 x *g* for 2 minutes to collect the droplets at the bottom of the tube, then the top 25 uL of dilute phase were pipetted off into a new tube. Then the tube containing the coacervate phase and dilute phase referred to as “BL” (bottom layer), and the top layer containing just the dilute phase referred to as “TL” (top layer) were placed directly into two separate scintillation tubes and scintillation counted. The partitioning of the radiolabeled RNA was calculated by first subtracting the background radiation from both the TL and BL. Then the counts from the TL were subtracted from the BL to give the counts for the coacervate phase only. Then the counts for the TL were multiplied by 2 because the dilute phase essentially is all of the 50 uL of reaction volume to provide the number of counts in the dilute phase. Then the total counts were calculated by adding the coacervate phase counts to the dilute phase * 2 counts. Then to calculate the fraction of RNA in coacervate phase, the coacervate phase counts are divided by the total counts 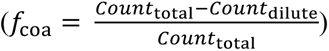.

### FRET of RNA duplex in coacervates

We used the methods published in literature by Nott et al.^56^ , Cakmak et al.^18^ and Choi et al. For Föster resonance energy transfer (FRET), ssRNA-Cy3 (5′-ACCUUGUUCC[Cy3]-3′) and its complementary sequence of Cy5-ssRNA (5′-[Cy5]GGAACAAGGU-3′) were used (same RNA sequences used in RNA partitioning experiments). Cy3 is the donor excited by 543 nm laser and its emission was collected between 555-625 nm. Cy5 is the acceptor excited by 633 nm laser and its emission was collected between 650 nm-750 nm. Three channels of fluorescence images were as following: *DD*_obs_ (Donor emission after donor excitation), *DA*_obs_ (Acceptor emission after donor excitation), *AA*_obs_ (Acceptor emission after acceptor excitation). The samples containing donor-only or acceptor-only dyes as stated in **Supplementary Table 4** were also imaged using same parameters of confocal microscope to get two correction values: *⍺* and *β* respectively; *⍺* = *DA*_donor_/*DD*_donor_, *β* = *DA*_acceptor_/*AA*_acceptor_. The corrected FRET efficiency (*E*_CT_) is calculated using following equation (1) to correct for the overlap of the emission and absorbance wavelengths of donor and acceptor dyes.

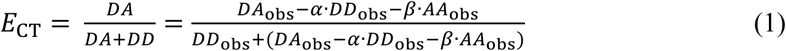

FRET efficiencies are calculated using the mean intensities from three samples prepared, and each sample has three images containing five droplets. FRET plots were processed from fluorescence intensity images using Fiji^69^ using averaged *⍺* and *β*. As controls, dsRNA in buffer and ssRNA in buffer were prepared to contain 5 µM of each RNA strands for enough fluorescence intensity by the same confocal microscope settings. Both low pH buffers (pH 4.8) and high pH buffer (pH 8.3) contain 10 mM phosphate, while salt concentrations are varied to be either low salt concentration (15 mM KCl and 0.5 mM MgCl_2_) or high salt concentration (600 mM KCl and 0.5 mM MgCl_2_ ).

---

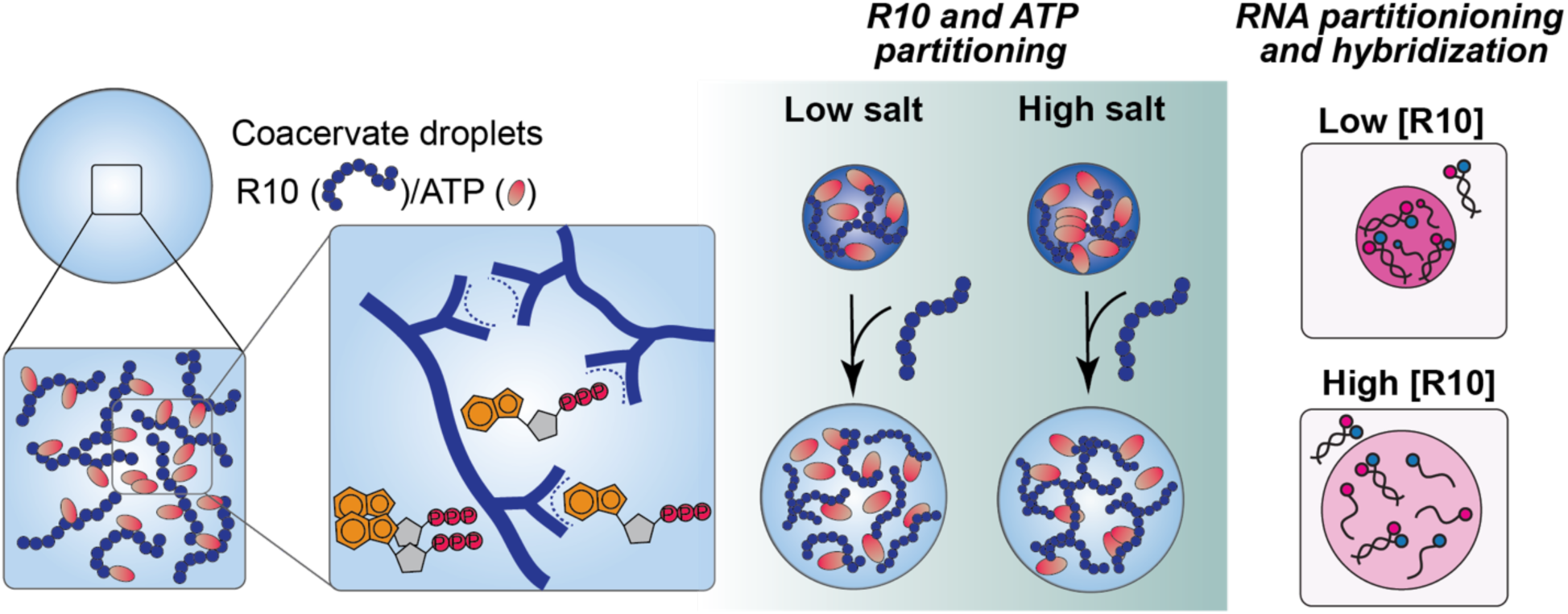

